# cGAS/STING-DEPENDENT SENSING OF ENDOGENOUS RNA

**DOI:** 10.1101/2022.05.16.492039

**Authors:** Tina Schumann, Santiago Costas Ramon, Nadja Schubert, Mohamad Aref Mayo, Melanie Hega, Servi-Remzi Ada, Lukas Sydow, Mona Hajikazemi, Yan Ge, Markus Badstübner, Stefan Lienenklaus, Barbara Utess, Lina Muhandes, Michael Haase, Luise Müller, Marc Schmitz, Thomas Gramberg, Nicolas Manel, Thomas Zillinger, Stefan Bauer, Alexander Gerbaulet, Katrin Paeschke, Axel Roers, Rayk Behrendt

**Author notes:** Correspondence should be addressed to R.B.

## Abstract

Defects in nucleic acid metabolizing enzymes lead to spontaneous but selective activation of either cGAS/STING or RIG-like receptor (RLR) signaling, causing a pathogenic type I interferon response and inflammatory diseases. In these pathophysiological conditions, cGAS-driven IFN production is linked to spontaneous DNA damage. Physiological, or tonic, IFN signaling on the other hand is essential to functionally prime nucleic acid sensing pathways. Here we show that low-level chronic DNA damage in mice lacking the Aicardi-Goutières syndrome gene *SAMHD1* reduced tumor-free survival when crossed to a p53-deficient, but not to DNA mismatch repair-deficient background. Increased DNA damage did not result in higher levels of type I interferon. Instead, we found that the chronic interferon response in SAMHD1-deficient mice was driven by the MDA5/MAVS pathway but required functional priming through the cGAS/STING pathway. Our work positions cGAS/STING upstream of tonic IFN signaling and highlights an important role of the pathway in physiological and pathophysiological innate immune priming.

**Summary:** Loss of the dNTPase and DNA repair enzyme SAMHD1 is associated with cancer and causes systemic autoimmunity. We show transformation-promoting spontaneous DNA damage and MDA5-driven but cGAS/STING-dependent chronic type I interferon production in SAMHD1-deficient mice.

## Introduction

Intracellular recognition of nucleic acids is essential in antiviral and in anti-tumor immunity, but uncontrolled activation of this machinery in the context of severe viral infections and tissue damage can cause detrimental inflammation. The same potent and therefore dangerous response is initiated when cells fail to control emergence of endogenous nucleic acids in amounts that exceed normal physiological levels or that lack secondary modifications, which would prevent autoactivation of nucleic acid receptors such as the dsDNA sensor cGAS or the RIG-like dsRNA receptors (RLR) RIG-I, MDA5 and LGP2 (Ablasser and Hur, 2020). RIG-I senses blunt-end dsRNA based on the presence or absence of 5’modifications, while MDA5 requires dsRNA stem structures of so far unknown minimal length (Ablasser and Hur, 2020). The enzyme cGAS senses nucleosome-free dsDNA inside of cells and produces the second messenger 2’3’cGAMP, a direct ligand for the cyclic-di-nucleotide sensor STING (de Oliveira Mann and Hopfner, 2021). Nucleosome-free dsDNA has been shown to accumulate in cells after DNA damage and therefore DNA damage has been identified as a primary pathogenic event in an increasing number of sterile inflammatory conditions that are driven by cGAS/STING-dependent cytokine production (Crow and Stetson, 2021). Both, activated STING and MAVS, recruit the kinase TBK1 leading to downstream activation of type I interferon (IFN) and NF-κB responses. IFN in turn acts autocrine and paracrine to stimulate expression of primarily antiviral genes via the type I IFN receptor (IFNAR). In many, but not in all cell types, expression of several pattern recognition receptors, including cGAS and the RLRs is regulated via this positive-feedback loop and depends on tonic IFN signaling, which leads to severely reduced PRR levels in IFNAR-deficient cells compared to IFNAR-competent cells (Behrendt et al., 2013; Schaupp et al., 2020). This results in a broad antiviral immune-defect in mice (Cervantes-Barragán et al., 2009; Schaupp et al., 2020) and in humans with inborn errors of the IFN system (Zhang et al., 2020). How exactly tonic IFN signaling is established and how this impacts physiological and pathophysiological immune priming is an emerging topic. Aicardi-Goutières syndrome (AGS) is a monogenic systemic autoimmune disease that is associated with high levels IFN in peripheral blood and in cerebrospinal fluid (Rodero and Crow, 2016). Mutations in AGS genes lead to spontaneous but selective activation of either RIG-like-receptor or cGAS/STING signaling (Crow and Stetson, 2021). The latter pathway has been implicated in the pathogenesis of AGS with underlying defects in the gene *SAMHD1* (AGS5) (Rice et al., 2009, 1; Maelfait et al., 2016; Daddacha et al., 2017; Coquel et al., 2018). SAMHD1 has at least two different functions: It’s an enzyme with deoxynucleoside triphosphate triphosphohydrolase (dNTPase) activity (Goldstone et al., 2011). Through this activity, SAMHD1 limits the availability of dNTPs in resting cells, which hinders replication of pathogens like retroviruses that depend on cellular dNTP supply (Hrecka et al., 2011; Laguette et al., 2011). Furthermore, increased levels of SAMHD1 in relapsed hematopoietic tumors have been shown to degrade nucleotide analogues thereby diminishing the efficacy of chemotherapy (Schneider et al., 2017; Herold et al., 2017). Moreover, SAMHD1-deficient tumor cells can be selectively killed through targeting the nucleotide metabolism (Davenne et al., 2020) making it an attractive anti-cancer drug target. A second function of SAMHD1 is reflected by the recruitment of the enzyme to sites of DNA double strand breaks (DSB) and to stalled replication forks. There, it interacts with the endonuclease CtIP and with the MRN complex to facilitate end resection in preparation for DSB repair by homologous recombination and to enable fork-restart (Daddacha et al., 2017; Coquel et al., 2018). The latter function does not require dNTPase activity and is mediated by the C-terminal region of SAMHD1. Failure in recruiting the DNA repair machinery by SAMHD1 results in spontaneous DNA damage and in the release of self-DNA that has been suggested to activate cGAS (Daddacha et al., 2017; Coquel et al., 2018). In agreement with its functions in DNA repair, SAMHD1-deficient patient fibroblasts showed a spontaneous transcriptional signature of interferon-stimulated genes (ISGs) and increased numbers of DNA double strand breaks (Kretschmer et al., 2015). The latter caused a chronic activation of the p53 pathway and senescence (Kretschmer et al., 2015). Impaired DNA repair predisposes to malignant transformation and consequently, mutations in SAMHD1 have been identified in many different tumors (Clifford et al., 2014; Rentoft et al., 2016). However, mutations found in cancer cells scatter across the whole *SAMHD1* gene (reviewed by (Mauney and Hollis, 2018) and do not allow for general conclusion about a definitive mechanism that could explain how the protein prevents malignant transformation. Here, two scenarios seem plausible: Loss of SAMHD1 dNTPase activity could affect the composition of cellular dNTP pools, which has a direct effect on the fidelity of replicative polymerases and could cause a mutator phenotype.

This has been widely studied in cancers originating from de-regulation of the ribonucleotide reductase complex (Aye et al., 2015). On the other hand, loss of SAMHD1-mediated DNA repair activity could cause increased numbers of DSBs and delay their repair, which might promote the selection of cell clones that inactivated cell cycle checkpoints to overcome this block. However, none of these scenarios have been experimentally addressed using in vivo models.

In addition to these established functions of SAMHD1, one group reported an exoribonuclease activity of the protein (Choi et al., 2015; Ryoo et al., 2014, 2016). However, RNAse activity of SAMHD1 was not reproduced by other studies (Seamon et al., 2015; Antonucci et al., 2016; Bloch et al., 2017; Yu et al., 2021), including our own (Wittmann et al., 2015). Therefore, if and how SAMHD1 regulates RNA metabolism in cells, remains to be fully elucidated.

In contrast to patients, loss of SAMHD1 in mice caused a mild activation of the type I IFN system but no systemic autoimmunity (Behrendt et al., 2013; Rehwinkel et al., 2013; Thientosapol et al., 2018). The IFN response was shown to be meditated via the cGAS/STING pathway (Maelfait et al., 2016). Furthermore, as opposed to studies in human cells lacking SAMHD1, no spontaneous DNA damage and no increased frequency of spontaneous tumors have been described in three independently generated SAMHD1-deficient mouse strains (Behrendt et al., 2013; Rehwinkel et al., 2013; Thientosapol et al., 2018). The lack of detectable DNA damage but the presence of a spontaneous IFN response in these mutant mice is an unresolved incoherence with the current understanding of how IFN is induced in SAMHD1-deficient cells.

Here we show low-level chronic DNA damage in SAMHD1-deficient mice that is detected by the p53 pathway. We found that inactivation of SAMHD1 in p53-deficient mice, but not in mice with defective DNA mismatch repair, reduced the tumor-free survival. Surprisingly, increased DNA damage did not amplify the spontaneous IFN response in SAMHD1-deficient mice. In contrast, we found that IFN is induced via the RNA sensor MDA5. Using SAMHD1-deficient mice as a model, we show that innate immune sensing of endogenous RNA through the RLR pathway requires functional priming via the cGAS/STING pathway.

## Results

### 1. Low level chronic DNA damage in SAMHD1-deficient mice

We and others previously reported a mild spontaneous IFN response in *Samhd1* knockout mice, which was dependent on the cGAS/STING pathway, suggesting that it was triggered by endogenous DNA (Behrendt et al., 2013; Maelfait et al., 2016). So far, however, there were no reports about spontaneous DNA damage in SAMHD1-deficient mice, which led us to ask if IFN in these mice is induced by an alternative mechanism to the human or if evidence of spontaneous DNA damage has been overlooked. Indeed, gene set enrichment analysis (GSEA) of whole transcriptome data from peritoneal macrophages revealed that only two types of pathways, reflecting an ongoing inflammatory response, including type I IFN, and replication stress were enriched in SAMHD1-deficient macrophages over control macrophages (Fig. 1A). This is in line with previous reports about human cells and suggests that also mouse SAMHD1 acts on stalled replication forks and in DNA repair (Daddacha et al., 2017; Coquel et al., 2018). Furthermore, nuclei of primary SAMHD1-deficient MEFs showed slightly elevated levels of γH2AX, a genuine marker for DNA strand breaks, when compared to littermate control MEFs (Fig 1B). Of note, in our hands this difference equilibrated after passage four and was no longer detectable in clones that overcame replicative senescence (not shown). DNA damage in erythroid precursors results in DNA double strand breaks and the rapid emergence of micronucleated reticulocytes followed by an increase of micronucleated erythrocytes, which can act as a short and long-term memory of genotoxic insults, respectively. After sub-lethal whole body irradiation, frequencies of micronucleated reticulocytes were increased by 4-fold in irradiated vs. non-irradiated *Samhd1*^+/+^ mice (Fig. 1C). This increase was doubled in *Samhd1^Δ/Δ^* mice, indicating higher susceptibility of SAMHD1-deficient mice to genotoxic stress (Fig. 1C). Next, we compared the steady-state frequencies of micronucleated erythrocytes in the peripheral blood of several SAMHD1-deficient mouse strains in our colony. Compared to the respective SAMHD1-proficient control of the same mutant strain, animals that lacked SAMHD1 consistently showed higher frequencies of micronucleated erythrocytes in peripheral blood, indicative of low-level chronic spontaneous DNA damage in these mice (Fig. 1D). Taken together, our results suggest that in SAMHD1-deficient mice genome replication and DNA repair are impaired resulting in low levels of chronic DNA damage.

**Figure 1:**
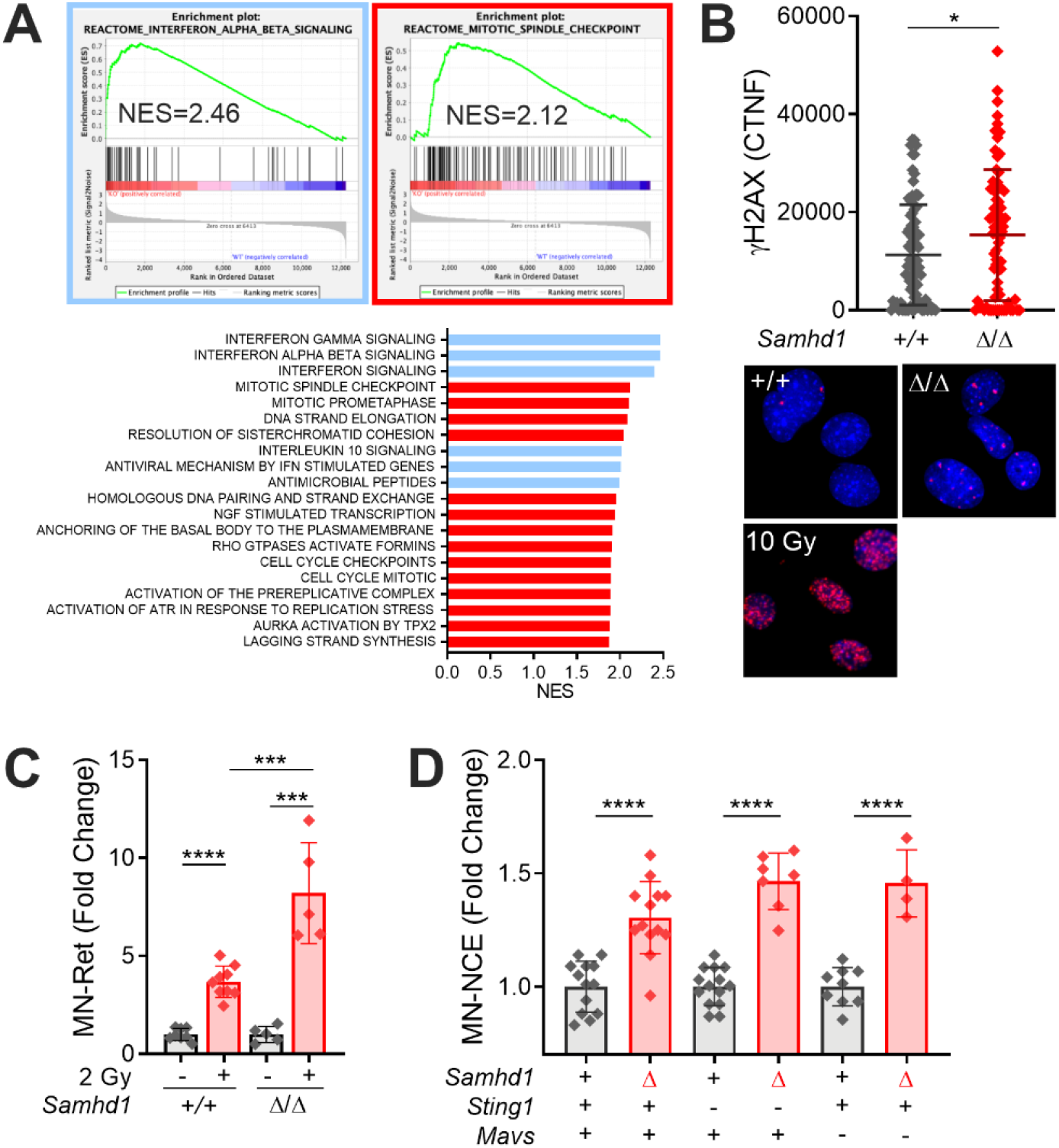
Low-level chronic DNA damage in SAMHD1-deficient mice. (A) Gene set enrichment analysis against the Reactome gene set collection (MSigDB) showing that exclusively gene sets of immune pathways (blue) and DNA replication (red) are enriched in *Samhd1^Δ/Δ^* vs. *Samhd1^+/+^* peritoneal macrophages. (B) Corrected total nuclear fluorescence (CTNF) of the γH2AX signal in pre-senescent primary MEFs of the indicated genotypes (Student’s t test) and representative immunofluorescence pictures. 10 Gy = positive control, analyzed 60 min after irradiation. (C) Change in micronucleated reticulocytes (MN-Ret) before (-) and 48 hrs after (+) whole body gamma-irradiation with a dose of 2 Gy in *Samhd1^+/+^* (n=8) and *Samhd1^Δ/Δ^* (n=5) mice. Fold change compared to mean of *Samhd1^+/+^* before irradiation is shown (One-way ANOVA followed by Tukey’s multiple comparison test). (D) Relative change in micronucleated normochromatic erythrocytes (MN-NCE) from peripheral blood of mice with the indicated genotypes. Fold change was calculated for each genetic background between *Samhd1^+/+^* (+) and *Samhd1^Δ/Δ^* (Δ). For *Mavs* and *Sting1*: + = *WT/WT*, - = *KO/KO*, n≥4 for each group (Student’s t test). *=p<0.05, ***=p<0.001, ****=p<0.0001.

### 2. Loss of SAMHD1 reduces tumor-free survival of mice lacking p53, but not of mice with defective DNA mismatch repair

Next, we asked why SAMHD1-deficient mice do not develop increased frequencies of spontaneous tumors although they showed spontaneous DNA damage in various cell types. Patient fibroblasts lacking SAMHD1 activate the p53 pathway in response to spontaneous DNA damage (Kretschmer et al., 2015). We reasoned that the low-level DNA damage in *Samhd1^Δ/Δ^* mice can be kept in check by p53-mediated damage responses and that inactivation of the p53 pathway might reveal how loss of SAMHD1 impacts genome stability in vivo. To address this question, we crossed SAMHD1-deficient mice to *Trp53^-/-^* mice, which predominantly develop spontaneous thymic lymphoma (Jacks et al., 1994). In our colony, *Trp53^-/-^* mice showed a mean tumor-free survival of 28 weeks, which was reduced to 18 weeks in *Samhd1^Δ/Δ^Trp53^-/-^* mice (Fig. 2A). In a cohort of *Trp53^-/-^* mice sacrificed at 12 weeks of age we found slightly enlarged thymi compared to control mice. At the same age, *Samhd1^Δ/Δ^Trp53^-/-^* thymi were already significantly larger than thymi of the *Trp53^-/-^* group (Fig. 2B). Histopathologic examination of thymic sections from the same cohort revealed that in 5 out of 6 *Samhd1^Δ/Δ^Trp53^-/-^* mice the disease had already progressed to thymic lymphoma, while at that time point no lymphoma cells were identified in thymus sections from *Trp53^-/-^* mice (Fig. 2C & D). Subsequent immunophenotypic analysis by multicolor immunohistochemistry demonstrated that CD4^-^CD8^-^ double negative CD3^+^ T cells were the dominant population in thymi of *Samhd1^Δ/Δ^Trp53^-/-^* (Fig. 2E and S1) mice, and this population emerged in *Samhd1^Δ/Δ^Trp53^-/-^* mice before 12 weeks of age most evident in the CD25^-^ T cell subsets DN1 (Fig. 2F) and DN4 (Fig. S2A). Longitudinal quantification of T cell development further supported that the disease of *Trp53^-/-^* mice develops faster but not qualitatively different in the absence of SAMHD1 (Fig. S2A). PCR on total thymus DNA amplifying specific recombination events in the TCRβ genes (Martins et al., 2014) demonstrated T cell bi- or oligoclonality indicative of thymic T cell lymphoma (Fig. S2B). Our data thus show that additional loss of SAMHD1 accelerated malignant transformation in *Trp53^-/-^* mice most likely by enhancing DNA damage.

**Figure 2:**
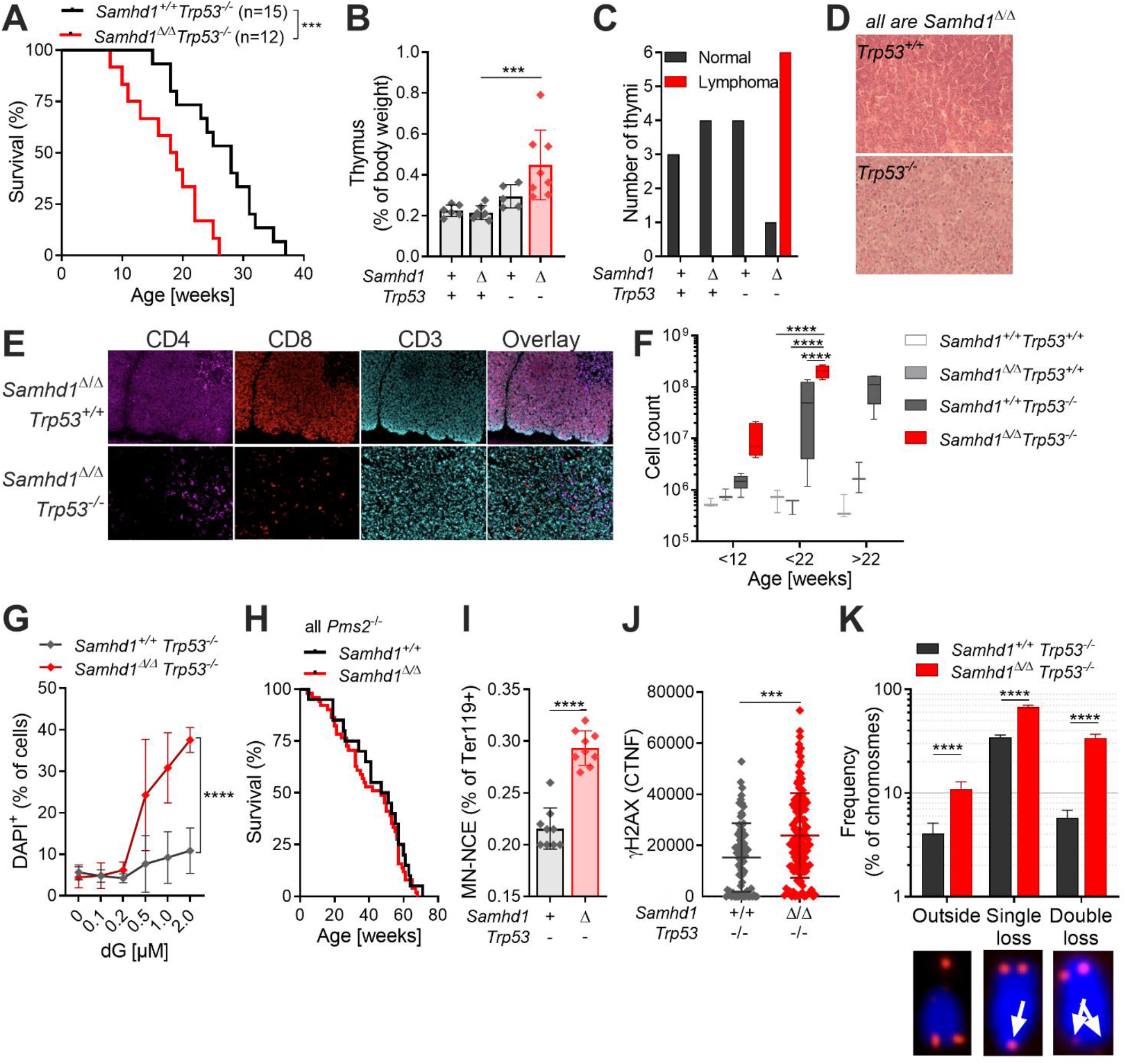
SAMHD1 prevents spontaneous DNA double strand breaks and accelerated transformation in of p53-deficient mice. (A) Tumor-free survival of *Samhd1^+/+^Trp53^-/-^* (n=15) and *Samhd1^Δ/Δ^Trp53^-/-^* (n=12) mice (log-rank test). For (B & C) + = *WT/WT*, - = *Δ/Δ* (B) Relative thymus weight of mice with the indicated genotypes at 12 weeks of age, n≥5 per group (One-way ANOVA followed by Sidak’s multiple comparison test). (C) Thymi of 12 weeks old mice with the indicated genotypes were examined for lymphoma formation by a trained histopathologist. Numbers of analyzed thymi in each group are shown and categorized according to the disease state. (D) Representative sections of a normal *Samhd1^Δ/Δ^Trp53^+/+^* (upper) and a *Samhd1^Δ/Δ^Trp53^-/-^* lymphoma bearing thymus. Sections were stained with H&E. (E) Representative multicolor immunohistochemistry staining for T cell lineage markers of thymic sections from mice with the indicated genotypes. See also Figure S1. (F) Cell counts of CD4^-^CD8^-^CD44^+^CD25^-^ DN1 immature T cells over time in the thymus of mice with the indicated genotypes. Complete dataset of T cell development in Figure S2, n≥3 for each group and time point (Two-way ANOVA followed by Tukey’s multiple comparison test). (G) Survival of *Samhd1^Δ/Δ^Trp53^-/-^* (n=3) and of *Samhd1^+/+^Trp53^-/-^* (n=3) immortalized thymic fibroblasts after treatment with deoxy guanosin (dG) at the indicated concentrations for 48 h. Representative of two independent experiments is shown (Two-way ANOVA). (H) Tumor-free survival of *Samhd1^Δ/Δ^Pms2^-/-^* (n=51) and of *Samhd1^+/+^Pms2^-/-^* mice (n=20). (I) Frequency of micronucleated normochromatic erythrocytes (MN-NCE) in peripheral blood of *Samhd1^Δ/Δ^Trp53^-/-^* (n=9) and of *Samhd1^+/+^Trp53^-/-^* (n=9) (Student’s t test). (J) Corrected total nuclear fluorescence (CTNF) of the γH2AX signal in pre-senescent primary MEFs from *Samhd1^Δ/Δ^Trp53^-/-^* and *Samhd1^+/+^Trp53^-/-^* mice. Representative result of two independent experiments is shown (Student’s t test). (K) Telomer integrity was quantified by FISH in 20 metaphases of immortalized thymic fibroblasts from *Samhd1^Δ/Δ^Trp53^-/-^* and from *Samhd1^+/+^Trp53^-/-^* mice (Student’s t test). ***=p<0.001, ****=p<0.0001.

We then asked if the altered nucleotide metabolism might contribute to malignant transformation in *Samhd1^Δ/Δ^Trp53^-/-^* mice. Due to altered dNTP levels, tumor cells lacking SAMHD1 can be selectively killed by 2’-deoxy-guanosin (dG) (Davenne et al., 2020). Indeed, we observed that immortalized *Samhd1^Δ/Δ^Trp53^-/-^* but not *Trp53^-/-^* thymic fibroblasts were hyper-sensitive to dG treatment, confirming that aberrant nucleotide metabolism caused by deficiency for SAMHD1 in cancer cells is an attractive drug target (Fig. 2G). Imbalanced dNTP levels decrease the fidelity of replicative polymerases and increase numbers of DNA mismatch mutations (Aye et al., 2015). In such a scenario, loss of SAMHD1 would be expected to reduce the tumor-free survival of DNA mismatch repair (MMR)-deficient mice. To test this hypothesis, we crossed *Samhd1^Δ/Δ^* mice to *Pms2^-/-^* mice, which lack functional MMR and develop spontaneous lymphoma (Baker et al., 1995). Surprisingly, and in contrast to our observations in *Trp53^-/-^* mice, loss of SAMHD1 did not significantly reduce the tumor-free survival of *Pms2^-/-^* mice (Fig 2H; 50% mean survival *Samhd1^Δ/Δ^Pms2^-/-^* 47 weeks, *Pms2^-/-^* 49 weeks, log-rank p=0.4052). This suggested that loss of SAMHD1 in mice is not associated with a strong mutator phenotype and that accelerated transformation seen in *Trp53^-/-^* mice lacking SAMHD1 is mainly driven by other forms of DNA damage.

To better understand the molecular events leading to reduced survival of *Samhd1^Δ/Δ^Trp53^-/-^* mice, we quantified the frequency of micronucleated erythrocytes in peripheral blood and found higher frequencies in *Samhd1^Δ/Δ^Trp53^-/-^* mice compared with *Trp53^-/-^* mice (Fig. 2I). In line with these observations, we detected a higher γH2AX signal in *Samhd1^Δ/Δ^Trp53^-/-^* versus *Trp53^-/-^* primary MEFs further supporting overall increased spontaneous DNA damage inflicted by additional loss of SAMHD1 in *SAMHD1^Δ/Δ^* in *Trp53^-/-^* mice (Fig. 2J). Independently of its role as a dNTPase, SAMHD1 recruits the MRN complex to promote homologous recombination at sites of DSBs and to re-start stalled replication forks. Telomeres consist of repetitive DNA sequences and form R-loop structures, both of which can lead to replication fork stalling. In cells with a defective shelterin complex, which protects telomeres from being recognized by the DNA repair machinery, SAMHD1 has been shown to prevent telomere breakage and the formation of extrachromosomal (“outsider”) telomere signals (Majerska et al., 2018). Quantification of telomere integrity in transformed *Samhd1^Δ/Δ^Trp53^-/-^* versus Trp53^-/-^ thymic fibroblasts by FISH revealed higher frequencies of chromosomes displaying single or double telomere loss or the characteristic outsider telomere signals (Fig. 2K). Our data suggests that loss of SAMHD1 in mice inflicts spontaneous DNA damage that is counteracted by a p53 response and in the absence of p53 accelerates tumor development. Accelerated transformation is more likely to be the result of increased numbers of DSBs in “difficult-to-replicate” regions, like telomeres, rather than being caused by a pronounced mutator phenotype.

### 3. Additional DNA damage does not amplify the spontaneous IFN response in SAMHD1-defcient mice

Cancer cell-intrinsic activation of the cGAS/STING pathway restricts tumor growth. This can be achieved by exogenous stimulation using synthetic ligands (reviewed in (Demaria et al., 2019)) or by promoting unphysiological accumulation of endogenous nucleic acids (Vanpouille-Box et al., 2019; Ishizuka et al., 2019). To investigate the role of endogenous DNA sensing in controlling tumor development in *Samhd1^Δ/Δ^Trp53^-/-^* mice, we crossed this mouseline to a STING-deficient background (Goldenticket mouse, *Sting1^GT/GT^*). Loss of STING had no effect on tumor-free survival of *Samhd1^Δ/Δ^Trp53^-/-^* mice (Fig. 3A) suggesting that STING signaling is not crucial in controlling the growth of *Samhd1^Δ/Δ^Trp53^-/-^*-deficient tumors. We and others previously reported that loss of p53 increased DNA damage and potentiated the spontaneous cGAS/STING-dependent IFN response in mice (Hiller et al., 2018) and in iPSCs (Giordano et al., 2022) with a defect in the essential DNA repair enzyme RNaseH2, strongly implicating genome damage in the generation of immune stimulatory DNA in this model. To better understand the lack of an effect of STING-deficiency in our *Samhd1^Δ/Δ^Trp53^-/-^* mouse model, we quantified the spontaneous activation of the type I IFN system. We observed increased levels of the surface ISG Sca-1 on peripheral blood lymphocytes (Fig. 3B) and increased transcription of several ISGs in peripheral blood of *Samhd1^Δ/Δ^* vs. *Samhd1^+/+^* control mice (Fig. 3C). However, this response was not further increased by additional loss of p53 (Fig. 3B & C). We made similar observations in bone marrow-derived macrophages from *Samhd1^Δ/Δ^Trp53^-/-^* and control mice (Fig. 3D). Of note, like in the p53 model, ISG transcription was similar between *Samhd1^Δ/Δ^* and *Samhd1^Δ/Δ^ Pms2^-/-^* mice (Fig. 3E).

**Figure 3:**
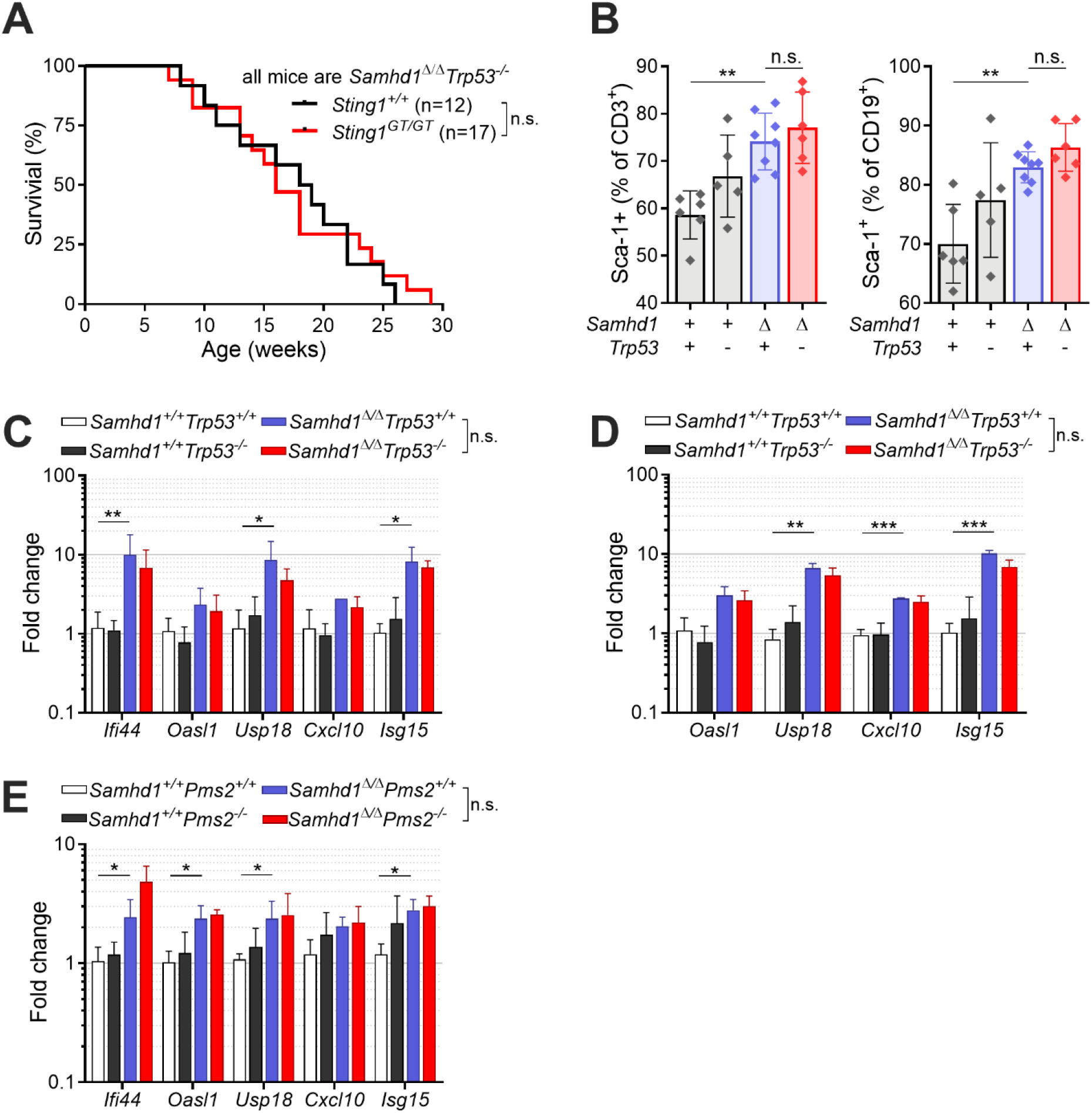
Redundant role of STING signaling in controlling tumor growth and IFN production in *Samhd1^Δ/Δ^Trp53^-/-^* compared with *Samhd1^+/+^Trp53^-/-^* mice. (A) Tumor-free survival of *Samhd1^Δ/Δ^Trp53^-/-^* on a STING-deficient (*Sting1^GT/GT^*) and STING-proficient (*Sting1^+/+^*) genetic background (log-rank test). (B) Frequency of DAPI^-^Sca-1^+^CD3^+^ T cells (left) and DAPI^-^Sca-1^+^CD19^+^ B cells (right) in peripheral blood of mice with the indicated genotypes (One-way ANOVA followed by Tukey’s multiple comparison test). (C – E). Relative transcript levels of ISGs in peripheral blood (C and E) and BMDMs (D) of mice with the indicated genotypes. Fold change compared to the mean of *Samhd1^+/+^Trp53^+/+^* (C and D) or *Samhd1^+/+^Pms2^+/+^* (E) are shown, n=3 for each group in each experiment (multiple t tests were performed). *=p<0.05, **=p<0.01, ***=p<0.001, n.s. = not significant.

In contrast to our previous observations in RNaseH2-deficient mice (Hiller et al., 2018), and despite higher levels of spontaneous DNA damage upon loss of SAMHD1 in p53-deficient mice (Fig. 2), this did not result in a stronger activation of the IFN system rendering STING signaling irrelevant in the control of tumor growth. We conclude that the p53-dependent DNA damage response does not function to prevent the generation of IFN-inducing endogenous nucleic acids in SAMHD1-deficient cells.

### 4. MDA5 drives IFN production in a cGAS/STING-dependent manner in *Samhd1^Δ/Δ^* mice

Our observation that increased DNA damage did not boost the IFN response in SAMHD1-deficient mice led us to investigate the exact role of cGAS/STING signaling in this model. To address whether STING signaling is important in SAMHD1-deficient mice, we treated *Samhd1^Δ/Δ^* mice for two weeks with 10 mg/kg of the STING antagonist H-151 (Haag et al., 2018). Pharmacologic inhibition of STING was able to reduce the transcription of ISGs in peripheral blood, demonstrating that STING is required for spontaneous IFN production in SAMHD1-deficient mice and that it represents a valuable therapeutic target to treat inflammatory conditions ensuing from defects in SAMHD1 (Fig. S3A). Until now, spontaneous IFN production caused by bi-allelic mutations in any of the AGS-related genes could be explained by activation of either the cGAS/STING pathway or the RLR pathway. In order to confirm that loss of SAMHD1 selectively activated the cGAS/STING pathway, we turned to a genetic approach to directly compare the relevance of intracellular DNA and RNA sensing in SAMHD1-deficient mice. As expected from our data with the STING inhibitor, knockout of *Sting1* completely blunted the ISG response in *Samhd1^Δ/Δ^* mice (Fig. 4A & B). To our surprise, also loss of MAVS abrogated the ISG response in SAMHD1-deficient peritoneal macrophages. This suggested that in contrast to mutations in other AGS enzymes, both, intact STING and MAVS signaling are required for the spontaneous IFN production in SAMHD1-deficient mice (Fig. 4A & B). Next, we asked if the nucleic acid sensors upstream of MAVS and STING were chronically activated in SAMHD1-deficient cells, or if there was a direct cross-talk between the two pathways at the level of STING and MAVS. To this end, we used post-replicative senescence *Samhd1^Δ/Δ^* MEFs hat retained a spontaneous ISG response, which could be rescued by lentiviral expression of murine SAMHD1 (Fig. S3B). In these MEFs we inactivated cGAS, RIG-I (*Ddx58*) and MDA5 (*Ifih1*) using CRISPR/Cas9 (Fig. S3C). As observed before, knockout of *cGas* completely blunted the transcription of several ISGs in SAMHD1-deficient cells (Fig. 4C). For the RIG-like receptors, only loss of MDA5 but not of RIG-I was able to reduce the mRNA levels of the ISGs tested (Fig. 4C). To further substantiate our finding that the spontaneous ISG response is indeed MDA5-dependent, we crossed *Samhd1^Δ/Δ^* mice to *Ifih1^-/-^* mice and analyzed the transcriptome of peritoneal macrophages. Like in previous experiments, inflammatory pathways and pathways indicating DNA replication stress were enriched in peritoneal macrophages from *Samhd1^Δ/Δ^* mice compared with control mice (Fig. 4D & 4E). In contrast, in peritoneal macrophages from *Samhd1^Δ/Δ^ Ifih1^-/-^* mice only pathways related to DNA replication and cell cycle progression remained enriched, when compared to control macrophages (Fig. 4D & 4E). The absence of upregulated inflammatory pathways in *Samhd1^Δ/Δ^ Ifih1^-/-^* mice indicated that MDA5 is chronically activated in SAMHD1-deficient mice and suggested the presence of endogenous immune-stimulatory dsRNA.

**Figure 4:**
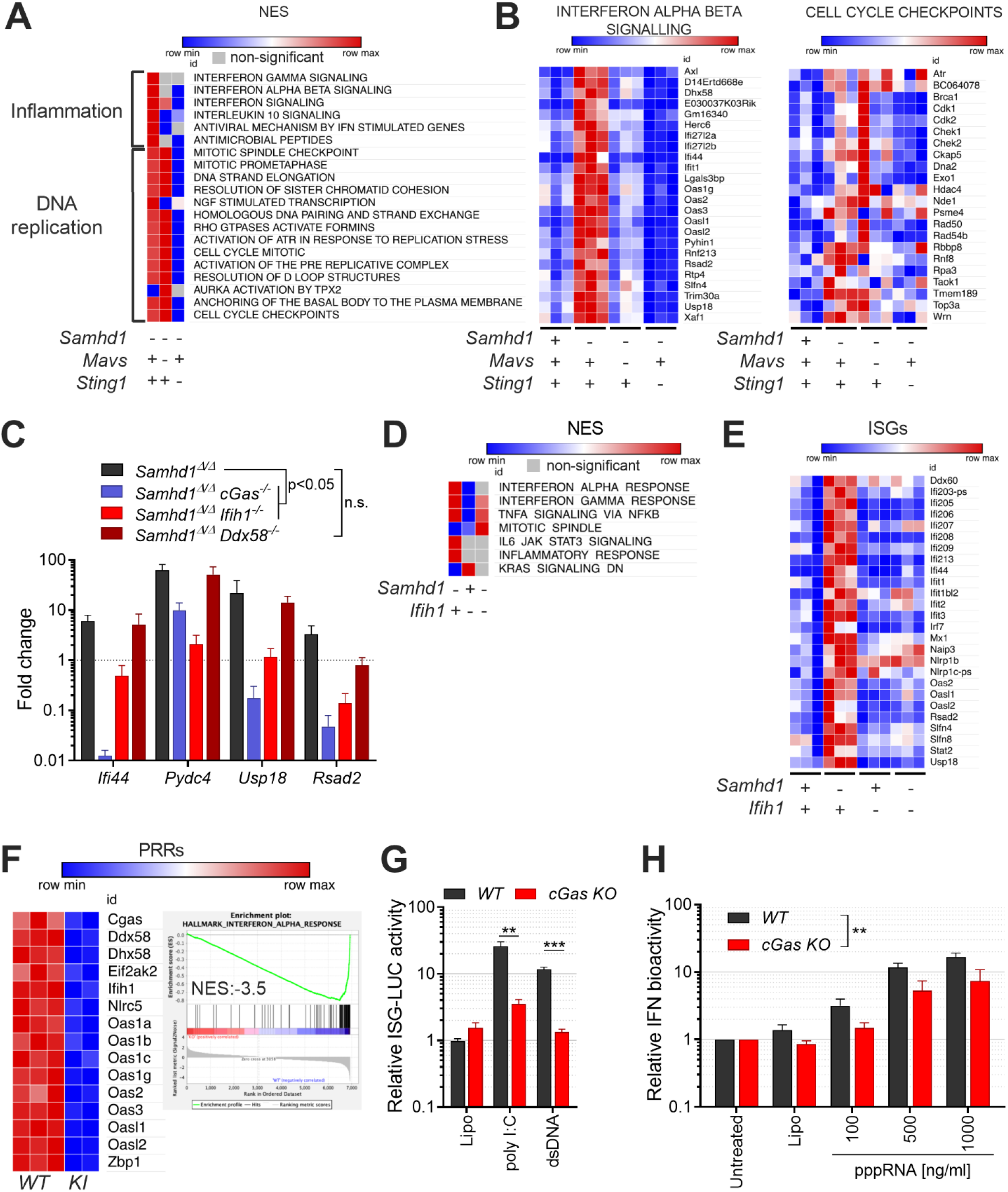
MDA5 drives spontaneous IFN production in a cGAS/STING-dependent manner in *Samhd1^Δ/Δ^* mice. For the whole figure - = homozygous null, + = homozygous wild type. (A) Enrichment of Reactome gene sets (MSigDB) in the transcriptome of peritoneal macrophages from mutant mice compared with littermate wild type controls of *Samhd1^Δ/Δ^* mice. (B) Normalized read counts for the indicated ISG transcripts (left) and transcripts of the CELL CYCLE CHECKPOINTS gene set (right) from the analysis shown in (A). (C) Relative transcript levels of the indicated ISGs measured by qRT-PCR in post-replicative senescence *Samhd1^Δ/Δ^* MEFs with additional CRISPR-mediated inactivation of the genes *cGas* (n=4), *Ifih1* (n=3) and *Ddx58* (n=2). Data of two independent experiments were pooled and displayed as fold change compared to the mean of *Samhd1*^+/+^ MEFs (multiple t tests, summary of results is shown with p<0.05 as lowest significance level). (D) Enrichment of Reactome gene sets (MSigDB) in the transcriptome of peritoneal macrophages from *Samhd1^Δ/Δ^Ifih1^+/+^, Samhd1^+/+^Ifih1^-/-^* and *Samhd1^Δ/Δ^Ifih1^-/-^* compared with littermate *Samhd1^+/+^ Ifih1^+/+^control* mice. (E) Normalized read counts for the indicated ISG transcripts from the experiment shown in (D). (F) Normalized read counts for transcripts of pattern recognition receptors (PRRs) in BMDMs from *Samhd1^Δ/Δ^GFP-cGas^KI/KI^* (n=2) vs. *Samhd1^+/+^* (n=3) control mice and the enrichment plot for the gene set INTERFERON_ALPHA_RESPONSE of the Hallmark gene set (MSigDB). (G) Relative ISG-luciferase reporter activity in cGAS-competent (WT) and cGAS-deficient (cGas KO) LL171 cells 16 hours after lipofection with 100 ng/μl poly I:C and 1 μg/ml plasmid DNA (3 kb). Luciferase activity was normalized to the mean of Lipo-treated WT LL171 cells (Student’s t test). (H) BMDMs isolated from *cGas^+/+^* (WT) and *cGas^-/-^* (cGas KO) mice lipofected with 10 μg/ml plasmid DNA and the indicated amounts of pppRNA or incubated with 10 μg/ml DMXAA for 4 hours. Cells were washed and incubated for another 18 hours before type I IFN bioactivity in the supernatant was determined using LL171 ISG-LUC reporter cells (two-way ANOVA). *=p<0.05, **=p<0.01, ***=p<0.001.

We were still puzzled by the requirement for cGAS in this system that was independently observed by us and by other groups (Maelfait et al., 2016; Coquel et al., 2018). Our transcriptomes of STING-deficient peritoneal macrophages (Fig. 4B) and as well qPCR results of cGAS-deficient MEFs (Fig. 4C) showed that ISG-expression in the mutant cells is below the levels found in cGAS/STING-competent control cells. This suggested that cGAS primes the antiviral immune system, most likely by its sporadic activation in response to endogenous DNA. To address the relevance of cGAS-mediated immune priming in the absence of SAMHD1, we generated SAMHD1-deficient mice that express GFP-tagged cGAS (Gentili et al., 2019) from a hypomorphic allele reducing the cGAS protein level to around 20% compared to that in WT mice (Fig. S3D). Whole transcriptome sequencing of BMDMs from *Samhd1^Δ/Δ^GFP-cGas^KI/KI^* and control mice demonstrated reduced expression of ISGs including several pattern recognition receptors and among these RIG-I (*Ddx58*) and MDA5 (*Ifih1*) (Fig. 4F). Haploinsufficiency for *cGas* rescues mice lacking the DNase Trex1 from lethal autoimmunity (Gao et al., 2015; Gray et al., 2015), suggesting that even in terminally sick *Trex1^-/-^* mice, the amount of cGAS ligands is only slightly above the threshold of tolerance. In contrast to *Trex1^-/-^* mice, the spontaneous IFN response in SAMHD1-deficient mice is very weak as illustrated by the lack of a spontaneous signal in *Samhd1^Δ/Δ^* Ifnβ-LUC reporter mice (Fig. S3E) rendering it even more sensitive to fluctuations in the cGAS protein level. As reported for reduced levels of cGAS, reduced levels of RIG-I and MDA5 might de-sensitize the intracellular RNA sensing pathways. To test if the sensitivity of the RLR pathway is controlled by cGAS, we knocked out *cGas* in LL171 ISG-luciferase reporter cells. As expected, no luciferase activity was detected after transfection of plasmid DNA. However, also the response to transfected poly I:C was almost completely blunted (Fig. 4G). Next, we isolated BMDMs from *cGas^-/-^* and littermate control mice and transfected them with increasing amounts of 5’triphosphate dsRNA to stimulate a RIG-I response. As observed for the poly I:C response, the same amounts of RIG-I ligand only induced lower IFN levels in the supernatant of *cGas^-/-^* BMDMs compared with controls (Fig. 4H). Taken together, our results suggest that cGAS keeps expression of PRRs at functional levels, which enables sensing of endogenous dsRNA by MDA5 in cells lacking SAMHD1.

## Discussion

We showed that loss of SAMHD1 in mice leads to a DNA replication defect, which causes DNA damage that is counteracted by activation of a p53 response. Loss of SAMHD1 in tumor cells exacerbates DNA damage in difficult-to-replicate regions like telomeres, which might contribute to accelerated malignant transformation when DNA repair pathways are impaired. Our observations of increased telomere damage in tumor cells lacking SAMHD1 are in line with previous reports of SAMHD1 being part of the telomere proteome (Majerska et al., 2018; Lin et al., 2021) and with increased frequencies of R-loop structures in SAMHD1-defcient cells (Park et al., 2021). Contrary to our observations in the p53-deficient tumor model, loss of SAMHD1 in MMR-deficient mice had no effect on tumor-free survival. This was unexpected as earlier reports demonstrated that even *Samhd1^+/-^* mice had elevated dNTP levels and mutations in the ribonucleotide reductase complex cause a cancerogenic mutator phenotype as a results of altered dNTP levels (Rentoft et al., 2016; Aye et al., 2015). *Samhd1^Δ/Δ^* mice lack dNTPase and DNA repair activity suggesting that loss of neither activity leads to significant DNA damage that would be detected by the MMR machinery in vivo. Hence, we conclude that loss of SAMHD1 in mice does not cause a strong mutator phenotype.

Activation of the innate immune response has been shown to efficiently control tumor growth and various means for targeted activation of intracellular nucleic acid sensing pathways are currently developed to boost anti-tumor immunity (Demaria et al., 2019). As opposed to exogenous stimulation of the pathway, STING had no role in regulating the tumor-free survival of *Samhd1^Δ/Δ^Trp53^-/-^* mice suggesting that endogenous DNA damage in p53-deficient tumors only weakly, if at all, activates STING. The relevance of our observation is illustrated by the fact that every other tumor in humans carries homozygous inactivation of the p53 gene (Baugh et al., 2018) and supported by a report showing that cGAS, but not STING protects from malignant transformation in a chemical-induced model of colon cancer (Hu et al., 2021). In order to fully understand the differential roles of cGAS and STING in controlling tumor growth, dissecting their multifaceted roles in regulating anti-tumor immunity and in controlling DNA damage will be instrumental.

We recently reported evidence that the IFN response in *Trex1^-/-^* mice is linked to DNA replication (Schubert et al., 2022) and similar findings have been reported for *SAMHD1^-/-^* cells (Coquel et al., 2018). In both models, loss of p53 did not amplify the IFN response, while such p53-dependent amplification was observed in cells lacking RNaseH2, in which chromatin fragments activate cGAS (Hiller et al., 2018; Mackenzie et al., 2017; Giordano et al., 2022). This points to a differential involvement of the p53 pathway in the generation of immune stimulatory DNA as a result of DNA replication in cells lacking TREX1 or SAMHD1 compared with post-replicative DNA damage found in RNaseH2-deficient cells. Although in SAMHD1-deficient cells accumulation of ssDNA species and concomitant activation of cGAS has been reported, it still remains unclear whether these oligonucleotides represent direct ligands for the DNA sensor, which, under physiological conditions, is known to nucleate only in the presence of unprotected long dsDNA (Andreeva et al., 2017; Du and Chen, 2018). Thus, it remains possible that in SAMHD1-deficient cells cGAS activation had a different culprit. Furthermore, our results challenge a role of pathogenic DNA sensing in SAMHD1-deficient mice as inactivation of RLR sensing in SAMHD1-deficient but cGAS/STING-competent cells was sufficient to blunt the spontaneous IFN response. This cannot be explained by a lack of RLR-mediated immune priming because ISGs levels in MAVS-deficient mice were similar to that of wild type mice, while in the absence of STING they were below levels found in wild type mice (Fig. 4B). ISG transcription was also lower in *Samhd1^Δ/Δ^Sting1^GT/GT^* when compared to *Samhd1^Δ/Δ^Mavs^-/-^* mice (Fig. S3F).

We previously observed that pDCs are the main producers of tonic IFN in mice (Peschke et al., 2016), which was later shown to be induced in response to commensal bacteria activating TLR and MAVS signaling pathways (Schaupp et al., 2020). In the skin, microbiota induced de-repression of endogenous retroelements and a cGAS/STING-dependent IFN response, but in this study MAVS signaling was not investigated (Lima-Junior et al., 2021). Interestingly, in human macrophages phagocytosed gut commensal bacteria evoked an IFN response, which was co-dependent on STING and MAVS expression (Gutierrez-Merino et al., 2020), suggesting that innate immune priming in response to low-level chronic stimuli can be driven by innate sensing of endogenous DNA and RNA. Our transcriptome data indicated that in mouse peritoneal macrophages, in primary BMDMs and in the murine fibroblasts cell line LL171 the cGAS/STING pathway establishes tonic IFN signaling and baseline expression of antiviral genes, including RLRs. This places cGAS/STING signaling upstream of RLR sensing in these cell types, because the absence of this pathway leads to impaired cytoplasmic RNA sensing (Fig. 4 G & 4H). Similar findings have been reported in the context of RNA virus infections (Schoggins et al., 2014; Parker et al., 2018). As the IFN response in SAMHD1-deficient mice is weak, we propose that loss of cGAS/STING signaling in cells lacking SAMHD1 de-sensitizes the RLR pathways and increases tolerance against RNA ligands, thereby preventing spontaneous induction of IFN despite the presence of an endogenous MDA5 ligand. To this end, dsRNA originating from endogenous retroelements has been shown to activate MDA5 in cells lacking the AGS gene ADAR1 (Ahmad et al., 2018) and after DNA damage induced by chemotherapy (Clapes et al., 2021). De-repression of endogenous retroelements is not only a physiological response (Lima-Junior et al., 2021; Young et al., 2012; Yu et al., 2012) but it is also a general stress-response (Simon et al., 2019; De Cecco et al., 2019), and SAMHD1-deficient cells display signs of replication stress including spontaneous DNA damage as shown here and previously by other groups (Daddacha et al., 2017; Coquel et al., 2018). Therefore, it is tempting to speculate that the stress response alone is sufficient to promote aberrant transcription and processing of endogenous RNA from, but not limited to, the vast numbers of retroelement loci, which might lead to autorecognition by RNA sensors in SAMHD1-deficient cells.

Taken together, our work suggests that in SAMHD1-deficient cells endogenous dsRNA represents the primary nucleic acid ligand that drives IFN production and implicates an important role of the cGAS/STING pathway in physiological and pathophysiological innate immune priming.

## Material and Methods

### Mice

*Samhd1^Δ/Δ^* (Behrendt et al., 2013), *cGas^-/-^* (Schoggins et al., 2014), *Mavs^-/-^* (Michallet et al., 2008), *Sting1^GT/GT^* (Sauer et al., 2011), *Ifih1^-/-^* (Gitlin et al., 2006), *Trp53^-/-^* (Jacks et al., 1994), *Pms2^-/-^* (Baker et al., 1995), *Trex1^-/-^* (Morita et al., 2004), *ΔβLUC^KI/KI^* (Lienenklaus et al., 2009) and *GFP-cGas^KI/KI^* (Gentili et al., 2019) mice were described previously. *Trp53^-/-^* (#002101) and *Pms2^-/-^* (#010945) were purchased from The Jackson Laboratory. Mice were housed under specific pathogen-free conditions at the Experimental Center of the University of Technology Dresden. All animal experiments were done according to institutional guidelines on animal welfare and were approved by the Landesdirektion Sachsen (11-1/2010-33, 24-1/2013-12, 24/2017; 88/2017).

### Mouse embryonic fibroblasts

MEFs were generated by standard procedures. In brief, E11.5 mouse embryos were dissected and decapitated. After removal of internal organs, tissue was cut into small pieces, digested with 1x trypsin (0.25 %, Invitrogen) for 30 min at 37°C and disaggregated by pipetting. The cell suspension was cultured in DMEM (Gibco) supplemented with 10 % heat-inactivated fetal calf serum, 100 U/ml Penicillin, 100 mg/ml Streptomycin, 1x non-nonessential amino acids (all Biochrom) and 100 μM β-mercaptoethanol (Gibco). After 24 h, non-digested tissue aggregates were removed and the cells were kept cultivated in complete DMEM medium at 37°C and 5 % CO2 under atmospheric oxygen.

### Thymic fibroblasts

Thymi were homogenized and passed through a 40 μm cell filter. The single cell suspension was cultured in RPMI 1640 medium supplemented with 10 % heat-inactivated fetal calf serum, 100 U/ml Penicillin, 100 mg/ml Streptomycin, 1 mM sodium pyruvate and 2 mM L-Alanyl L-glutamine (all Biochrom). Surviving cells were kept cultivated in complete RPMI 1640 medium at 37°C and 5 % CO_2_ under atmospheric oxygen.

### In Vitro Differentiation of BMDMs

Bone marrow cells were cultured over night in RPMI 1640 medium supplemented with 10 % heat-inactivated fetal calf serum, 100 U/ml Penicillin, 100 mg/ml Streptomycin, 1 mM sodium pyruvate and 2 mM L-Alanyl L-glutamine (all Biochrom). The next day, non-adherent cells were transferred to new dishes and differentiated for six days in RPMI medium (supplemented as described) containing 30 % L929 supernatant. An equal amount of fresh differentiation medium was added after three days. Four days later, the attached cells were harvested, counted and seeded in RPMI + 15 % L929 supernatant to perform the experiments. Cells were cultured at 37°C and 5 % CO_2_ under atmospheric oxygen.

### Micronucleus flow assay

Retrobulbar blood sample was mixed with heparin/PBS (250 units/ml, Biochrom) and fixed by adding 20 μl of the mixture to 2 ml ice-cold methanol, inverted and stored at −80 °C. Quantification of micronucleated erythrocytes was performed as described previously (Balmus et al., 2015). Briefly, fixed blood cells were washed in bicarbonate buffer and stained using antibodies against CD71 (Southern Biotech, #1720-02, 1:200) and Ter119 (eBioscience, #48-5921-82, 1:500) in the presence of 1μg/μl RNase A. After washing and addition of 1 μg/ml PI, cells were analyzed by flow cytometry and gated for single Ter119^+^ CD71^+^ PI^+^ micronucleated reticulocytes (MN-Ret) and Ter119^+^ CD71^-^ PI^+^ micronucleated normochromatic erythrocytes (MN-NCE).

### Transcriptomics of peritoneal macrophages

Peritoneal macrophages (DAPI^-^, CD11b^hi^, F4/80^hi^) were isolated by FACS using antibodies against CD11b (eBioscience, #11-0112, 1:1600) and F4/80 (Biolegend, #123114, 1:200), and total RNA was extracted using RNeasy Plus Mini Kit (Qiagen). For each experiment equal amounts of total RNA were used for poly-dT enrichment before library preparation and sequencing was performed as described before (Schubert et al., 2022). Reads were mapped to mouse genome GRCm39 followed by normalization, exploratory and differential expression analysis using DESeq2 (Love et al., 2014). Unless otherwise stated, DEG lists were generated by comparing all mutant mouse lines to the respective wild type group in each experiment and sorted in according to ascending padj. All transcripts with padj<0.05 were subjected to GSEA (Subramanian et al., 2005). To generate heatmaps all transcripts with padj<0.05 in the comparison Samhd1^Δ/Δ^ vs. Samhd1^+/+^ were extracted from lists containing normalized read counts of all genotypes in the respective experiment and the resulting sub list was displayed using Morpheus (https://software.broadinstitute.org/morpheus). New datasets are currently being deposited in the GEO database. Analysis in Figure 1 has been performed on a previously generated dataset GSE45358.

### Quantification of phosphorylated histone H2AX

25.000 MEFs were plated into 8-well chamber slides. Cells were left at 37°C over night to attach to the slide. After incubation, cells were washed in PBS, fixed in ice-cold methanol and washed again. For quantification of phosphorylated histone H2AX (γH2AX), slides were blocked at RT for 1 h in 1x Blocking reagent (Roche) and incubated at 4°C over night with a phospho-histone H2AX (pSer139) antibody (Cell Signaling Technology, #2577, 1:50). After washing with PBS, slides were incubated with a goat anti-rabbit-AF488 antibody (Thermo Fisher Scientific, #A-11034, 1:500) for 1h at RT in the dark. Slides were washed and nuclei were counterstained with 10 μg/ml DAPI in the mounting solution. Imaging was done on a Keyence fluorescence microscope and analyzed using ImageJ software (NIH). γH2AX foci in at least 50 fibroblast nuclei were counted. As positive control, wild type MEFs were gamma-irradiated with a dose of 10 Gy and analyzed 1 hour later.

### Histology

Thymi were formalin-fixed, paraffin-embedded, and cut into 3 μm sections. For H&E (haematoxylin-eosin) staining, sections were dyed with Mayer’s hemalum solution for 2 min, followed by staining with eosin and rinse with water for 30 seconds. The preparations were dehydrated again in an ascending alcohol series and washed in xylene. H&E sections were evaluated by a board-certified pathologist on a Zeiss Axioskop 2 microscope and photographs were made with an Axiocam 503 color camera using ZEN 2.5 (blue edition) software (Zeiss).

Multiplex immunohistochemical staining against CD3, CD4, and CD8 to assess T cell composition in thymi was performed on a Ventana Discovery Ultra Instrument. Briefly, antigen retrieval using cell conditioning 1 solution (Ventana Medical Systems) was performed at 95°C for 32 min, followed by incubation with the primary antibody against CD8 (eBioscience, #14-0195-82, 1:100) at 36°C for 32 min, the HRP-coupled secondary anti-rat OmniMap antibody (Ventana Medical Systems) for 12 min and finally Opal 520 fluorophore (Akoya Biosciences, 1:100) at RT for 8 min. Primary and secondary antibodies were removed by denaturation at 100°C for 24 min in cell conditioning 2 buffer (Ventana Medical Systems). The above described steps were repeated for CD4 (abcam, #ab183685, 1:500) with OmniMap anti-rabbit-HRP and Opal 570 fluorophore (Akoya Biosciences, 1:1000), and lastly CD3 (abcam, #ab16669, 1:50) with OmniMap anti-rabbit-HRP and Opal 690 fluorophore (Akoya Biosciences, 1:50). Finally, sections were counterstained with DAPI (Merck) and mounted with Fluoromount G mounting media (Southern Biotech). Sections were scanned at 100x magnification, regions of interest defined using Phenochart software (Akoya Biosciences), and multispectral images acquired at x200 magnification using the Ventra 3.0 Automated Imaging System (Akoya Biosciences). Upon spectral unmixing using inForm Software (Akoya Biosciences), images were exported and processed in ImageJ (NIH).

### Flow cytometry

Thymi were homogenized and passed through a 70 μm cell filter. Following washing with ice-cold FACS buffer (PBS/ 2 %FCS/ 2mM EDTA), cells were filtered again through a 70 μm cell filter. On samples from peripheral blood erythrocytes were lysed if leucocytes were analyzed. Cells were incubated with anti-CD16/CD32 (Biolegend, #101302, 1:200) at RT for 10 min to block Fc receptors and stained with the following antibodies in FACS buffer at 4 °C for 30 min: CD3e (eBioscience, # 11-0031, 1:200 or # 17-0031, 1:100), CD4 (eBioscience, #53-0041, 1:200), CD8a (eBioscience, #25-0081, 1:600), CD11b (eBioscience, #11-0112, 1:1600), CD19 (eBioscience, #25-0193, 1:200), CD25 (eBioscience, #12-0251, 1:800), CD44 (eBioscience, # 48-0441, 1:200), CD45R (B220) (eBioscience, # 47-0452, 1:100) and Ly-6A/E (Sca-1) (eBioscience, #17-5981, 1:200). After incubation, cells were washed and resuspended in FACS buffer. For dead cell exclusion, 1 μg/ml of PI was added to the cell suspension shortly before the analysis. Cells were analyzed using the FACSAria III (BD Bioscience) and evaluated with FlowJo Version 10 (Tree Star).

Peripheral blood was stained for Sca-1^+^ within the CD3^+^ and CD19^+^ populations.

### Telomere integrity

Quantification of telomere integrity was done with metaphase telomere Fluorescence In Situ Hybridization (FISH). Metaphase spreads were performed as previously published (Poon and Lansdorp, 2001). Briefly, the cells were cultured in 10 cm petri dishes and grown to 60% confluency. The cells were treated with 0.2μg/ml Colcemid (Merck #10 295 892 001) for 3 hours and incubated with hypotonic solution (75mM KCl). Swollen cells were washed with fixative solution (methanol:glacial acetic acid 3:1) and dropped on superfrost microscopic slides. Telomeres were stained with TelC-Alexa488 labelled PNA probe (Panagene, #F1004) as previously published (Awad et al., 2020). The slides were mounted with Fluoroshield mounting media containing DAPI (Sigma, #F6057-20ML) to stain the chromosomes. Images were acquired using a ZEISS Axio Observer microscope. The obtained images were analysed by evaluating the average telomere integrity per metaphase. Depending on the signal, the telomere phenotypes were categorised to fragile, outside, apposition and fusion.

### Quantitative RT-PCR

Total RNA was isolated using NucleoSpin RNA Kit (Macherey-Nagel) and reverse transcribed into cDNA using PrimeScript RT Reagent Kit (Takara) following the manufacturer’s instructions. Quantitative RT-PCR using Luna^®^ Universal qPCR Master Mix (New England BioLabs) was performed with the following cycling conditions on a CFX384 Touch Real-Time PCR Detection System (Bio-Rad): 10 min 95°C, 40 cycles of 95°C for 20 s, 60°C for 30 s. The used qRT-PCR primers are listed in Supplementary Table 1. Transcript levels were normalized to the housekeeping gene Tbp1. All samples were run in technical triplicates.

### CRISPR/Cas9 gene targeting in MEFs and LL171 cells

Cells were transfected with pSpCas9(BB)-2A-GFP (px458, Addgene) containing guide RNAs targeting genes *cGas, Ifih1* or *Ddx58*. Target sequences are given in Supplementary Table 1. Cells were selected with 3 μg/ml puromycin for 72 h and single cell clones were isolated in a 96-well format. Genotyping was performed by amplicon deep sequencing on a MiSeq using a protocol described by Lange et al., 2014, that was adapted to the target loci. Knock out of the target genes was determined genetically using the Outknocker tool (Schmid-Burgk et al., 2014) and functionally by the lack of response to specific ligands (Fig. S3).

### LL171 luciferase reporter assay

ISRE luciferase reporter expressing LL171 cells were cultured in DMEM (Gibco) supplemented with 10 % heat-inactivated fetal calf serum, 100 U/ml Penicillin, 100 mg/ml Streptomycin, 1x non-nonessential amino acids (all Biochrom) and 600 μg/ml G418. To analyze luciferase activity in cell supernatant, LL171 cells were seeded in supplemented DMEM w/o G418 in 96-well plates. Once cells are attached, DMEM was removed and IFN-containing cell supernatant was added over night to the cells. Luciferase assay was performed using the SpectraMax^®^ Glo Steady-Luc™ Reporter Assay Kit (Molecular Devices) according to the manufacturer’s instructions and relative luciferase activity was measured at the LUMIstar Omega (BMG Labtech) microplate reader.

### Western Blot

Cell pellets from *GFP-cGas^KI/KI^* and control BMDMs were lysed in 2x Laemmli buffer and incubated at 95 °C for 5 min. Proteins were separated on a 12 % denaturing acrylamide gel and subsequently transferred onto a nitrocellulose membrane (Amersham Hybond-ECL, GE Healthcare). Membrane was blocked using 1x Roti®Block (Carl Roth) for one hour and incubated over night at 4 °C with the primary antibodies against cGAS (Cell Signaling Technology, #31659, 1:1000), β-actin (Cell Signaling Technology, #4970, 1:10.000) and Cyclophilin B (Cell Signaling Technology, #43603, 1:20.000), diluted in 1x Roti®Block. Following washing with TBS/0.1% Tween (TBS-T), the membrane was incubated for one hour at RT with peroxidase-conjugated goat anti-rabbit secondary antibody (Cell Signaling Technology, #7074, 1:1000) and washed again. The Amersham ECL Prime Western Blotting Detection Reagent (GE Healthcare) was used for protein visualization using the Fusion FX (Vilber Lourmat) and Fusion FX7 Advanced imaging software. Signals were densitometrically analyzed using ImageJ software (NIH).

### Statistical Analysis

Data are shown as means ± SD. Statistical analysis was performed using GraphPad Prism 9. To compare the mean of two groups, Student’s t test (unpaired t test, two-tailed, 95% confidence intervals) was used. For the comparison of more groups one-way ANOVA or two-way ANOVA followed by either Tukey’s or Sidak’s multiple comparison test were used. Log-rank test was used to compare survival data. Significance levels in each figure are stated as follows: p ≤ 0.05, **p < 0.01, ***p < 0.001, ****p<0.0001.

---

**Supplementary Table 1.**
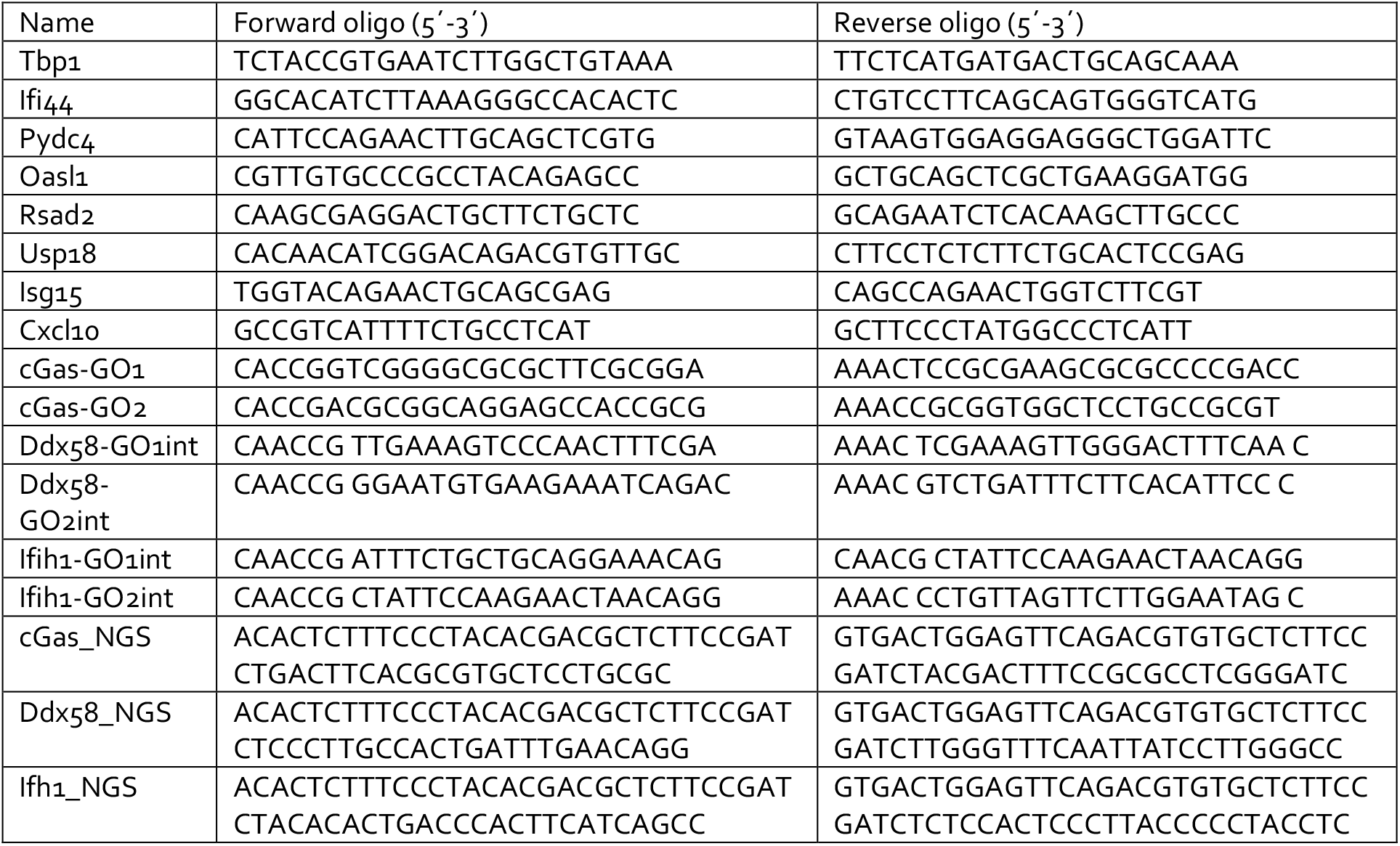
Oligonucleotides

## Author contributions

Conceptualization - R.B.; Methodology – R.B., S.C.R., T.S.;A.G., Li.M., Validation Verification - Li.M.,Me.H.; Formal Analysis R.B., A.G., M.A.M., Me.H, Y.G., S.R.A., R.O., N.S., Lu.M., Li.M., M.S.; Investigation - S.C.R., M.A.M., Y.G., S.R.A., R.O., L.S., N.S.,B.U., Lu.M., Li.M., M.S., A.G., R.B.; Resources - M.S., S.B., T.Z., N.M.; Data Curation – Y.G.; Writing – Original Draft - R.B., T.S., N.S.; Writing – Review & Editing - R.B., T.S., N.S., Visualization – R.B., N.S., M.H, Lu.M.; Supervision – R.B.,A.R, K.P., A.G.; Funding Acquisition – R.B., A.R., K.P.,S.B.

## Acknowledgements

R.B., A.R., K.P., S.B, T.Z. were supported by the DFG TRR/SFB237. R.B. was additionally supported by the DFG (BE-5877/2-1) and by an AGS Research Award. T.G. was supported by DFG 401821119/GRK 2504.

---

**Figure S1:**
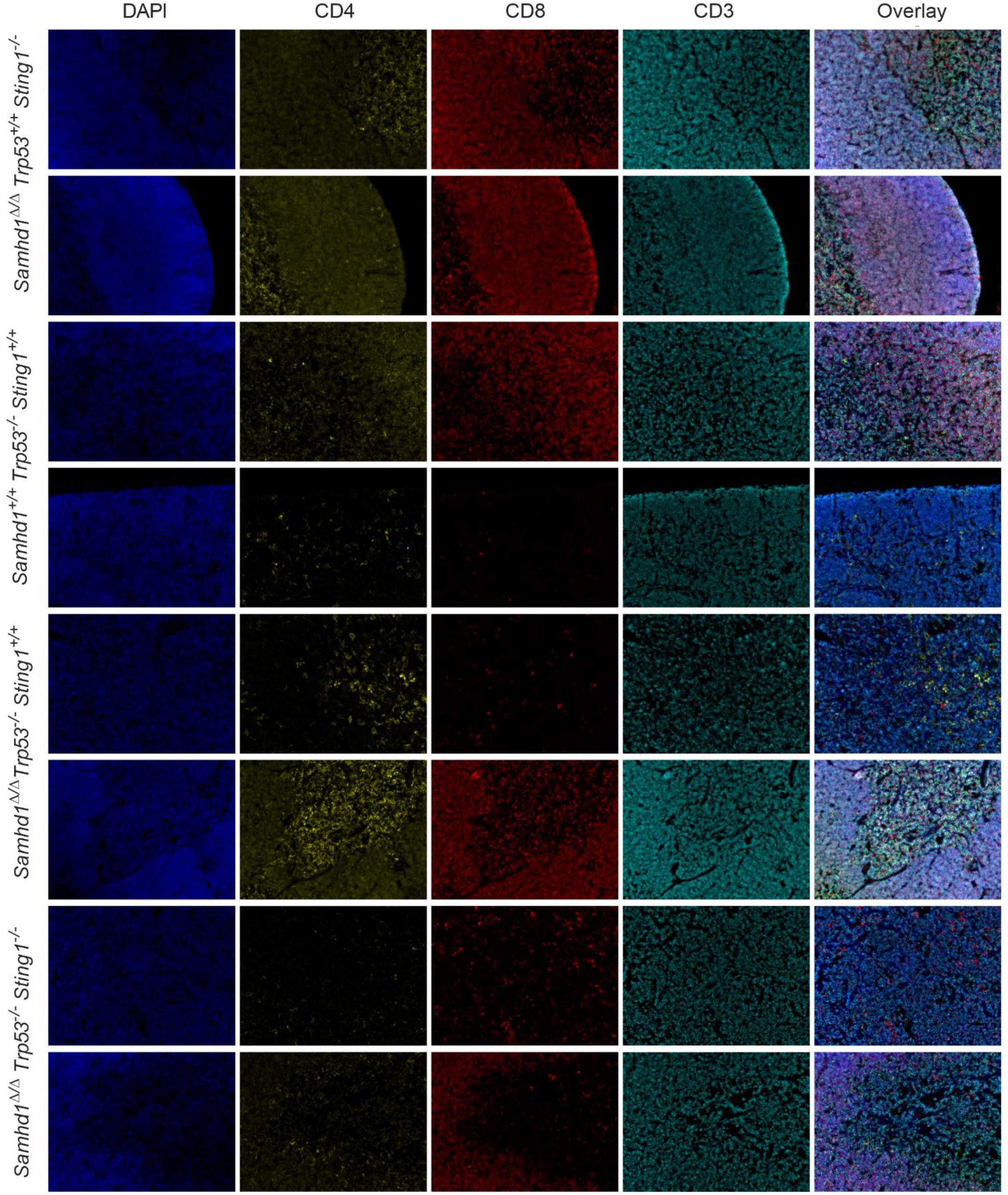
Characterization of T cell lineage marker expression by multicolor immunofluorescence in thymi of mice with the indicated genotypes. Related to Figure 2E.

**Figure S2:**
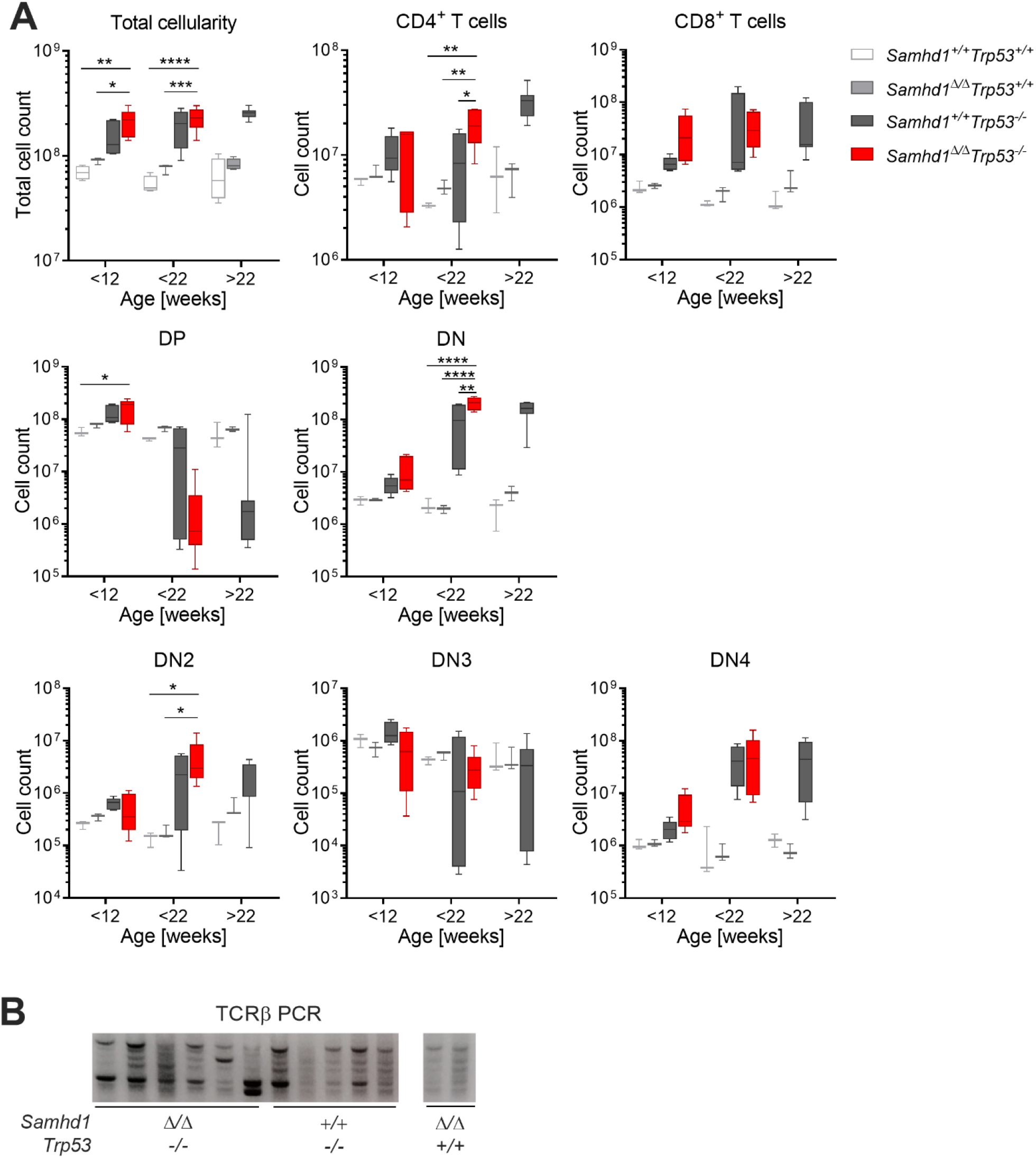
Aberrant T cell development in *Samhd1^Δ/Δ^Trp53^-/-^* and in *Samhd1^+/+^Trp53^-/-^* mice. Related to Figure 2. (A) Thymus parameters recorded by flow cytometry. Cells were gated based on scatter to exclude debris and DAPI-for living cells, before gating on the respective markers. DP= CD4^+^CD8^+^, DN=CD4^-^CD8^-^, DN1= CD4^-^CD8^-^CD44CD25^-^ (shown in Figure 2F), DN2= CD4^-^CD8^-^CD44^+^CD25^+^, DN3= CD4^-^CD8^-^CD44^-^CD25^+^, DN4= CD4^-^CD8^-^CD44^-^CD25^-^ (Two-way ANOVA followed by Tukey’s multiple comparison test). (B) DNA was extracted from total thymus of mice with the indicated genotypes. TCRβ loci were amplified by PCR using a combination of 22 primers binding in a V segment combined with one primer binding in J1.7. Similar results were obtained with primer J2.7 (not shown). Strategy according to (Martins et al., 2014). *=p<0.05, **=p<0.001, ***=p<0.001, ****=p<0.0001.

**Figure S3:**
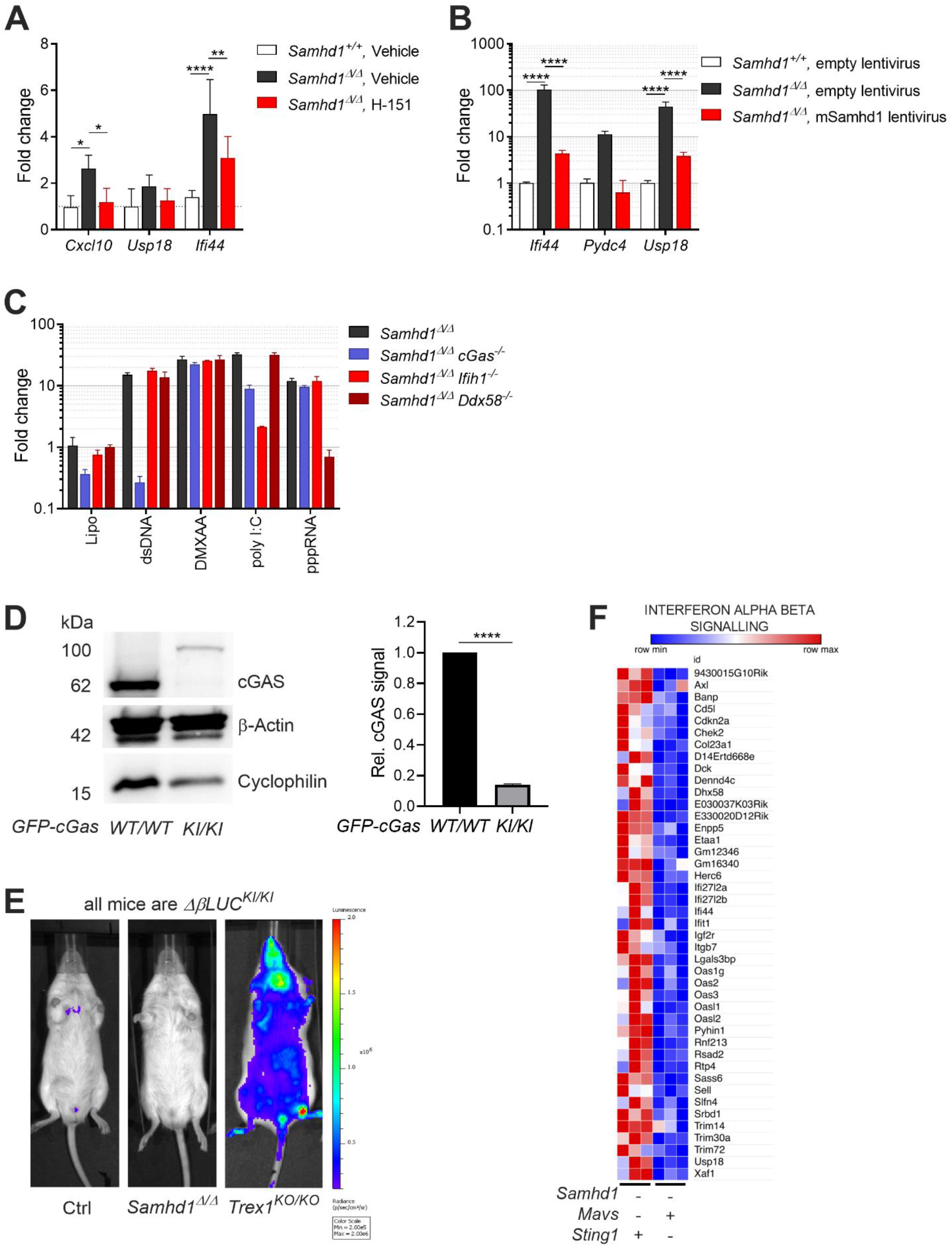
MDA5 drives spontaneous IFN production in a cGAS/STING-dependent manner in *Samhd1^Δ/Δ^* mice. Related to Figure 4. (A) *Samhd1^+/+^* and *Samhd1^Δ/Δ^* mice were treated i.p. with 10 mg/kg/day H-151 or vehicle for 14 days. Transcript levels of the indicated ISGs were determined in spleen. Fold change compared with the WT^-^vehicle group is shown, n=4 in each group (Two-way ANOVA followed by Tukey’s multiple comparison test). (B) Post-replicative senescence *Samhd1^Δ/Δ^ and Samhd1^+/+^* MEFs were transduced with empty lentivirus or a lentivirus which expresses the cDNA of murine *Samhd1* isoform1 as well as EYFP. Transduced cells were enriched by FACS for EYFP and transcript levels of the indicated ISGs were determined by qRT-PCR. Data of two independent measurements is displayed as fold change compared with the mean of *Samhd1^+/+^* MEFs transduced with empty lentivirus (Two-way ANOVA followed by Tukey’s multiple comparison test). (C) Relative transcript levels of the indicated ISGs measured by qRT-PCR in post-replicative senescence *Samhd1^Δ/Δ^* MEFs with additional CRISPR-mediated inactivation of the genes *cGas* (n=4), *Ifih1* (n=3) and *Ddx58* (n=2) after lipofection with 1 μg/ml plasmid DNA (dsDNA), 100 ng/ml poly I:C, 100 ng/ml pppRNA or incubation with 10 μg/ml DMXAA for 16 hours. Fold change compared to Lipo-treated *Samhd1^+/+^* MEFs is shown. (D) Representative western blot for cGAS in *GFP-cGas^KI/KI^* and *GFP-cGas^WT/WT^* control mice (left). Data from two independent experiments for densitometric quantification of cGAS signal relative to the signal for β-actin (right, Student’s t test). cGAS = 62 kDa, GFP-cGAS around 92 kDa. (E) Spontaneous *in vivo* Ifnb1-luciferase signal in *Samhd1^+/Δ^* (ctrl), *Samhd1^Δ/Δ^* and *Trexi^KO/KO^* mice. All mice were homozygous for the luciferase knock in (*ΔβLUC^KI/KI^*). (F) Normalized read counts of ISG transcripts in *Samhd1^Δ/Δ^Ifih1^-/-^ vs. Samhd1^Δ/Δ^Sting1^GT/GT^* mice. **=p<0.01, ****=p<0.0001.

